# Transcription factors overcome the repressive impact of Polycomb-associated methylation in tumors

**DOI:** 10.1101/2025.05.15.654174

**Authors:** Min-Kyeong Kwon, Goeun Park, Dayoung Go, Donghyun Park, Sridhar Hannenhalli, Sun Shim Choi

**Affiliations:** Division of Biomedical Convergence, College of Biomedical Science, Institute of Bioscience & Biotechnology, Kangwon National University, Chuncheon 24341, Korea; Planit Square Inc., Seoul 06242, Korea; Cancer Data Science Lab, Center for Cancer Research, National Cancer Institute, Bethesda, MD 20814

**Author notes:** Corresponding authors Sridhar Hannenhalli, Sun Shim Choi.

## Abstract

DNA methylation is a key epigenetic regulator often disrupted in cancer, yet how promoter methylation dynamics translate into transcriptional changes during cancer progression remains incompletely understood. Here, we employed targeted bisulfite sequencing and RNA-seq on paired tumor and non-tumor tissues from 80 Korean colorectal cancer (CRC) patients to map promoter methylation and gene expression dynamics. Promoters with high baseline methylation in non-tumor tissues tended to become hypomethylated in tumors, while those with low baseline methylation underwent partial hypermethylation. However, these changes did not consistently correlate with gene silencing or activation. Strikingly, promoters marked by Polycomb (PcG^+^) in non-tumor tissue were prone to hypermethylation yet often remained transcriptionally active in tumors, a paradox most prominent in transcription factor (TF) genes. In contrast, hypermethylation in PcG^−^ promoters was more consistently associated with transcriptional repression. Our findings suggest that epigenetic plasticity at PcG^+^ TF gene promoters can override the typically repressive effects of DNA methylation, potentially enabling tumors to maintain or enhance the expression of key regulatory genes. This highlights the importance of PcG occupancy in shaping the functional consequences of methylation changes during colorectal tumorigenesis, warranting deeper investigation into how these epigenetic adaptations drive cancer progression.

## Introduction

DNA methylation, the covalent addition of a methyl group to the 5-carbon of cytosine to form 5-methylcytosine (5mC), is a fundamental epigenetic modification that can heritably alter gene expression [1]. In mammals, this modification occurs predominantly at CpG dinucleotides, often within gene promoters and CpG islands (CGIs) [2, 3]. The prevailing model holds that promoter methylation suppresses transcription by either preventing transcription factor (TF) binding or by recruiting methyl-CpG–binding proteins (e.g., MeCP2) and histone deacetylases, leading to chromatin condensation [4-7]. Consequently, DNA methylation at promoters is generally linked to transcriptional repression, whereas unmethylated promoters are typically accessible and transcriptionally active.

Previous studies generally support this model. In many cancers, CGI promoter hypermethylation correlates with the silencing of tumor suppressor genes [8] —including those involved in DNA repair [9, 10], cell cycle regulation [11], and adhesion [12, 13] —thus contributing to tumorigenesis. In parallel, tumors exhibit global hypomethylation, particularly in repetitive sequences and intergenic regions, which is believed to promote chromosomal instability and aberrant gene activation [14, 15]. While CGI hypermethylation typically occurs at tumor suppressors, non-CGI promoters on the other hand, which are normally methylated in healthy cells, often become hypomethylated in cancer [16]. In some settings, this hypomethylation can activate genes (including oncogenes) [15].

Recent work however suggests a more complex relationship between promoter methylation and gene silencing; not all methylated promoters are silenced, and not all unmethylated promoters are active [17, 18]. One layer of complexity involves Polycomb group (PcG) proteins, which mark a large subset of developmental gene promoters with H3K27me3 [19]. These PcG-marked (PcG^+^) CGI promoters are typically unmethylated and yet transcriptionally repressed in normal cells [20]. PcG targeting thus represents an alternative repressive mechanism. However, during tumorigenesis, these loci are disproportionately targeted for *de novo* DNA methylation—originally assumed to “lock in” repression [20, 21]. Yet, studies now show that some PcG^+^ CGI genes can remain transcriptionally active despite promoter hypermethylation, especially if activated by distal enhancers or chromatin remodeling [22, 23]. Genes encoding TFs are of particular interest in this context, as many TF genes are PcG targets and play central roles in controlling cell identity and fate during tumorigenesis [24]. Certain TFs can even bind methylated DNA, suggesting a mechanism by which some genes may escape or even exploit methylation-induced silencing [25].

Most prior studies investigating the role of methylation on tumor-associated transcriptional changes have relied on comparing independent cohorts of tumor and normal tissue, potentially obscuring patient-specific regulatory dynamics [22, 26]. To address this specific limitation, we performed an integrative analysis of genome-wide promoter methylation and gene expression profiles in paired tumor and adjacent non-tumor tissues from 80 individuals with colorectal cancer (CRC). CRC is an ideal system for studying epigenetic disruption, as it is strongly associated with the CpG island methylator phenotype (CIMP) [11, 27]. In this study, we stratified gene promoters based on CpG content (CGI vs. non-CGI) and Polycomb state (PcG^+^ vs. PcG^−^), and we specifically investigated the dynamic changes in TF gene expression compared to their non-TF counterparts. By leveraging a paired-sample design, we directly assessed within-individual changes in DNA methylation and transcription, thereby revealing substantial inter-individual variability in methylation–expression relationships.

Our findings show that, although classical methylation patterns (CGI hypermethylation and non-CGI hypomethylation) hold, their effects on gene expression vary considerably among individuals. Most notably, PcG^+^ TF promoters frequently undergo hypermethylation in tumors yet paradoxically are often upregulated. These observations provide a more refined view epigenetic mechanism whereby TF genes may resist or subvert DNA methylation at their own promoters, enabling persistent or enhanced transcription in cancer. Overall, our study provides a nuanced view of gene regulation during tumorigenesis and suggests yet unknown alternative epigenetic mechanisms operating at PcG^+^ TF gene promoters as a potential hallmark of CRC and other cancers.

## Results

### Predominant CGI promoter hypermethylation and non-CGI promoter hypomethylation in CRC tumors

To investigate how DNA methylation in the promoter region affects gene expression during tumorigenesis, we analyzed targeted bisulfite sequencing (TBS) data for promoter-centered methylation and genome-wide transcriptome data (RNA-seq) from 80 Korean CRC patients. Each patient provided both non-tumor (NT) and tumor (T) samples, as described previously [28]. After quality control (QC), one outlier sample was removed, resulting in a final dataset of 79 matched NT-T pairs. We then examined the relationship between promoter methylation and gene expression for 14,042 protein-coding genes, of which 10,204 (72.7%) harbored CGIs in their promoters, and 3,838 (27.3%) did not (Figure 1A).

**Figure 1.**
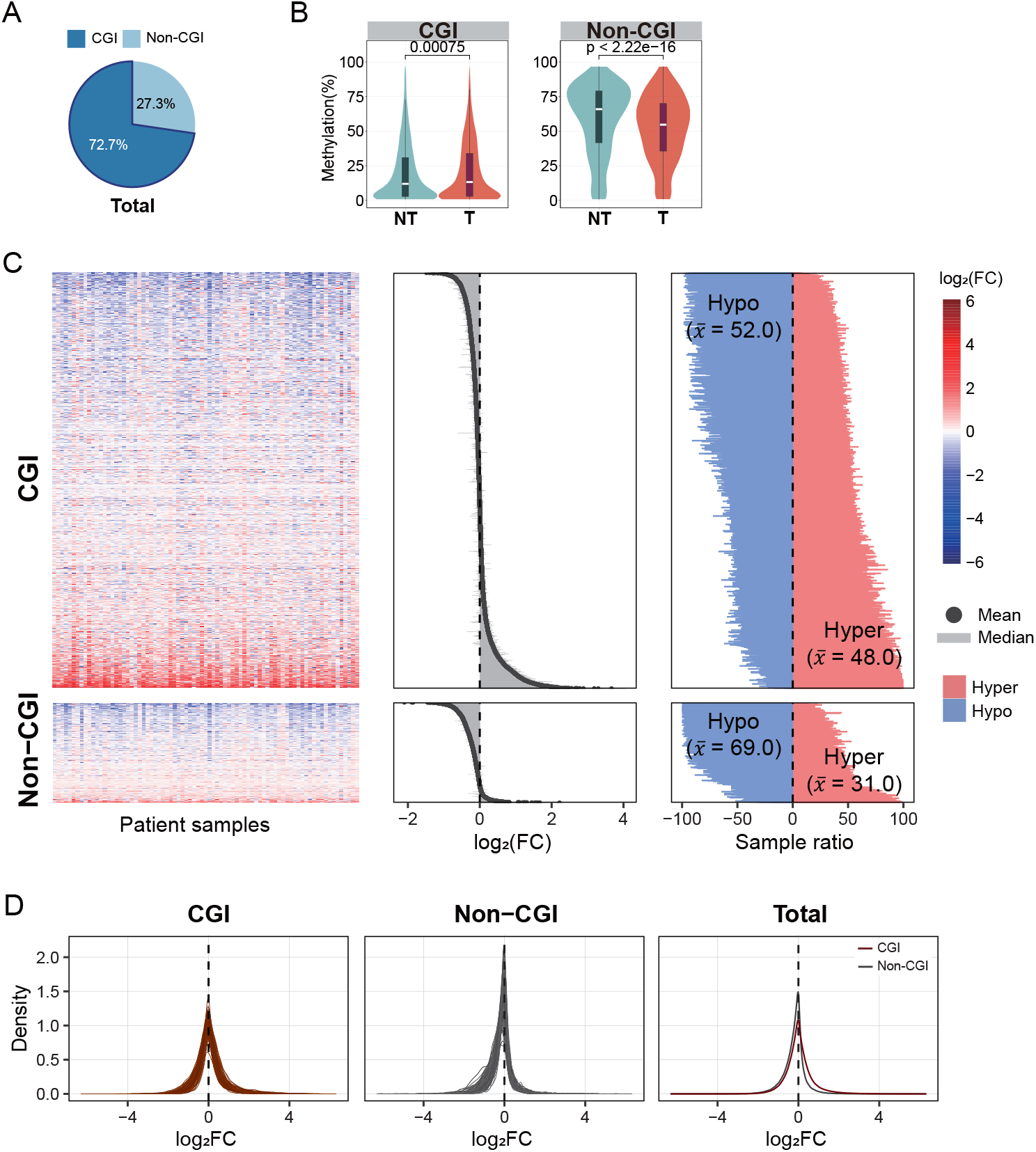
Overall landscape of promoter methylation in non-tumor vs tumor tissues. **A**. Pie chart showing the proportion of protein-coding genes with CGI vs. non-CGI promoters. **B**. Violin plots comparing promoter methylation levels (y-axis) between non-tumor (NT) and tumor (T) samples for CGI (left) and non-CGI (right) promoters. **C**. Heatmaps (left) showing gene-level methylation changes, measured as log_2_ FC of beta values between NT and T samples, across all patients. (Blue indicates hypomethylation, and red indicates hypermethylation). The middle figures show the median value as a light gray line and the average value as a dark gray dot for each gene. To the right, frequency plots illustrating how many patients exhibit hypermethylation or hypomethylation for each gene. The average frequency of hyper- or hypomethylation is indicated (x). CGI genes on top and non-CGI genes below. **D**. The density plot of log_2_FC values of methylation for each patient. The plot displays the distribution of log_2_FC in T compared to NT samples.

First, we evaluated the mean methylation level for each promoter region across all 79 pairs of samples. Consistent with the generally low methylation state of CGI promoters, they remained largely unmethylated in both NT and T tissues, yet, as a group, they exhibited a very modest but statistically significant increase in methylation (weak hypermethylation) in tumor samples (P = 0.00075). In contrast, non-CGI promoters—normally more methylated than CGI promoters—showed a marked decrease in methylation in tumor samples (strong hypomethylation; P < 2e-16) (Figure 1B).

Next, leveraging the paired data, we assessed methylation changes at the individual patient level by comparing matched NT and T samples. Figure 1C left panel shows for each gene promoter and each patient the tumor-vs-non-tumor methylation log-fold change (log_2_FC), revealing substantial inter-individual variability for both CGI and non-CGI genes; certain genes showed hypermethylation in approximately half of the samples and hypomethylation in the other half. As a result, their average log_2_FC was close to zero (middle panel of Figure 1C), indicating that many genes, regardless of promoter type, exhibited minimal overall methylation changes across the cohort. Interestingly, genes that were consistently hypermethylated across most samples were predominantly associated with CGI promoters, whereas those frequently hypomethylated were mainly linked to non-CGI promoters (right panel of Figure 1C). On average, 48% of CGI genes were more frequently hypermethylated, compared to only 31% of non-CGI genes; conversely, 69% of non-CGI genes were hypomethylated more frequently compared to 52% of CGI genes. Figure 1D, which represents cross-gene density plots of log_2_FC across individuals, supports the same conclusion as Figure 1C. These results broadly recapitulate previously observed hypermethylation at CGI promoters and hypomethylation at non-CGI promoters. Leveraging paired data, our analyses, while recapitulating the broad expected trends, reveals substantial heterogeneity and divergence across genes and patients, particularly at CGI promoters. Thus, it is of interest to further investigate instances of CGI promoter hypermethylation that exhibit consistency across patients, and features of such promoters.

### Methylation heterogeneity in CGI promoter genes in the context of Polycomb marks

Thus, we aimed to assess whether the methylation variability observed in CGI promoter-associated genes was related to Polycomb (PcG) complex occupancy. Toward this, we analyzed methylation patterns after dividing these genes into two groups: PcG^−^ (without PcG marks; 8,262 genes; 81%) and PcG^+^ (with PcG marks; 1,942 genes; 19%; Figure 2A). The presence or absence of PcG marks was determined using ChromHMM data for colon tissue from ENCODE (see Materials and Methods).

**Figure 2.**
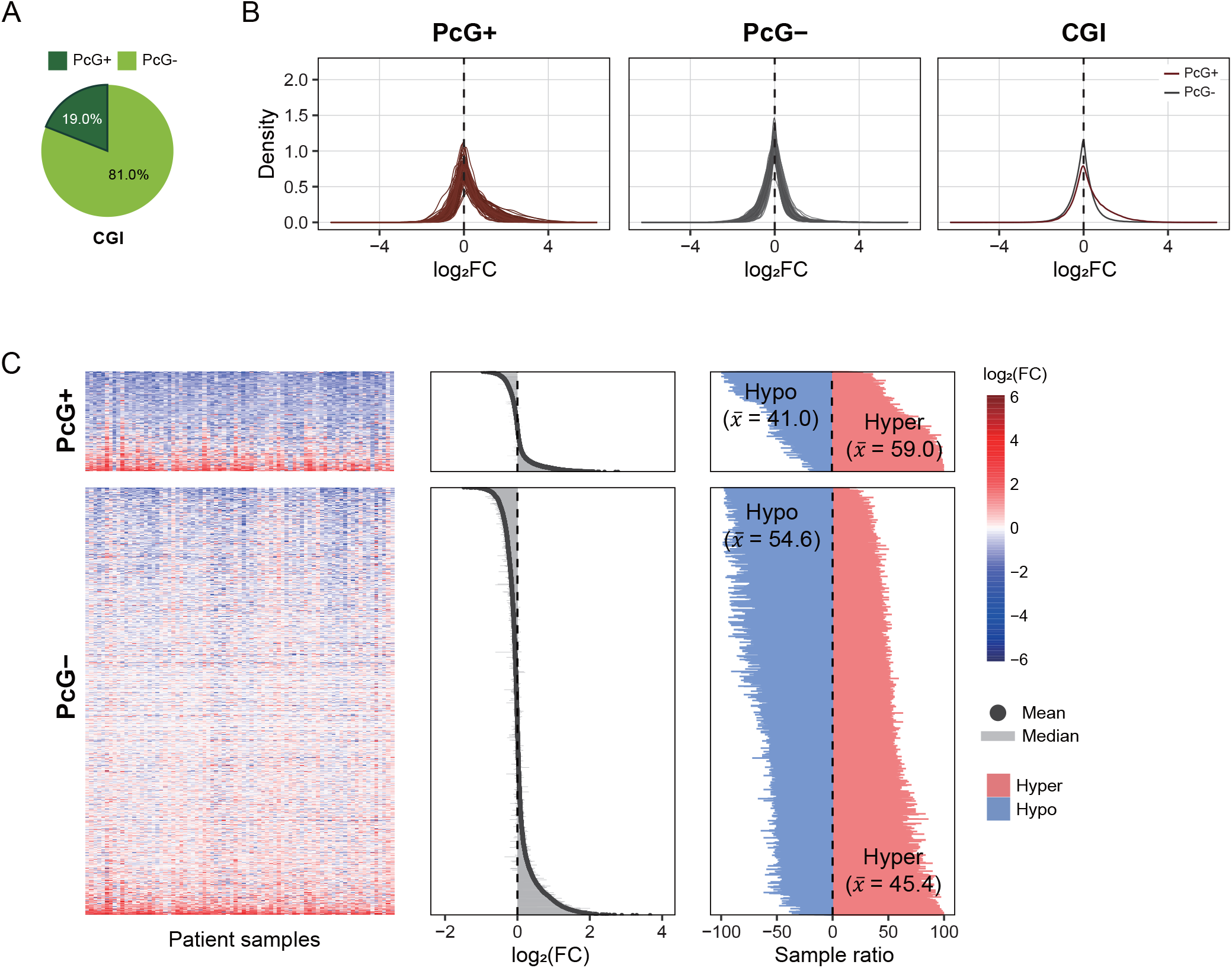
PcG marks and their effect in CGI methylation. **A**. Pie chart highlighting the fraction of CGI genes marked by PcG^+^ vs PcG^−^. **B**. The same analysis as in Figure 1D, the density plot of log_2_FC values of methylation for each patient **C**. The same analysis as in Figure 1C, PcG^+^ genes on top and PcG^−^ genes below.

We then performed analyses analogous to those in Figure 1, but separately in PcG-defined subgroup. As shown in Figure 2B, both PcG^+^ and PcG^−^ gene groups were centered around log_2_FC = 0, indicating little overall methylation change across individuals. However, the PcG^+^ group showed broader distribution tails, suggesting greater inter-individual variability. Most genes showed variable methylation patterns across samples rather than consistent hyper- or hypomethylation (Figure 2C, left and middle panels).

Notably, hypermethylation was more common among PcG^+^ genes (59%) than PcG^−^ genes (45.4%) (Figure 2C, right panel). This finding suggests that PcG-marked CGI promoters are preferential targets for hypermethylation in CRC, underscoring the importance of further investigating their role in gene regulation during tumorigenesis.

### Relationship between promoter methylation and gene expression across patients and promoter contexts is highly variable

To investigate the regulatory role of promoter methylation in gene expression, we analyzed their relationship across 79 matched CRC patient samples, with a particular focus on inter-individual variability and the dependence on PcG context.

As shown in Figure 3A, as expected, there is a clear inverse relationship between promoter methylation and gene expression in both CGI and non-CGI genes. This trend was consistent across both NT and T samples, supporting the canonical role of promoter methylation as a negative regulator. However, the shape and slope of the methylation–expression relationship varied between patients (Figure 3A), especially for non-CGI promoter genes having a broader dynamic range of methylation. Notably, in certain patients, this negative correlation appeared more pronounced in non-CGI promoters. Scatter plots of log_2_FC in promoter methylation versus gene expression further demonstrate this variability (Figure 3B). While most genes showed the canonical inverse relationship—hypermethylation with downregulation (hyper-dn) or hypomethylation with upregulation (hypo-up)—a substantial fraction displayed paradoxical patterns, with hypermethylated promoters linked to upregulation (hyper-up) or hypomethylated promoters linked to downregulation (hypo-dn). This suggests that when scrutinized based on paired dynamic data, the broadly presumed canonical relationship between promoter methylation changes and gene expression changes does not hold for a substantial fraction of genes.

**Figure 3.**
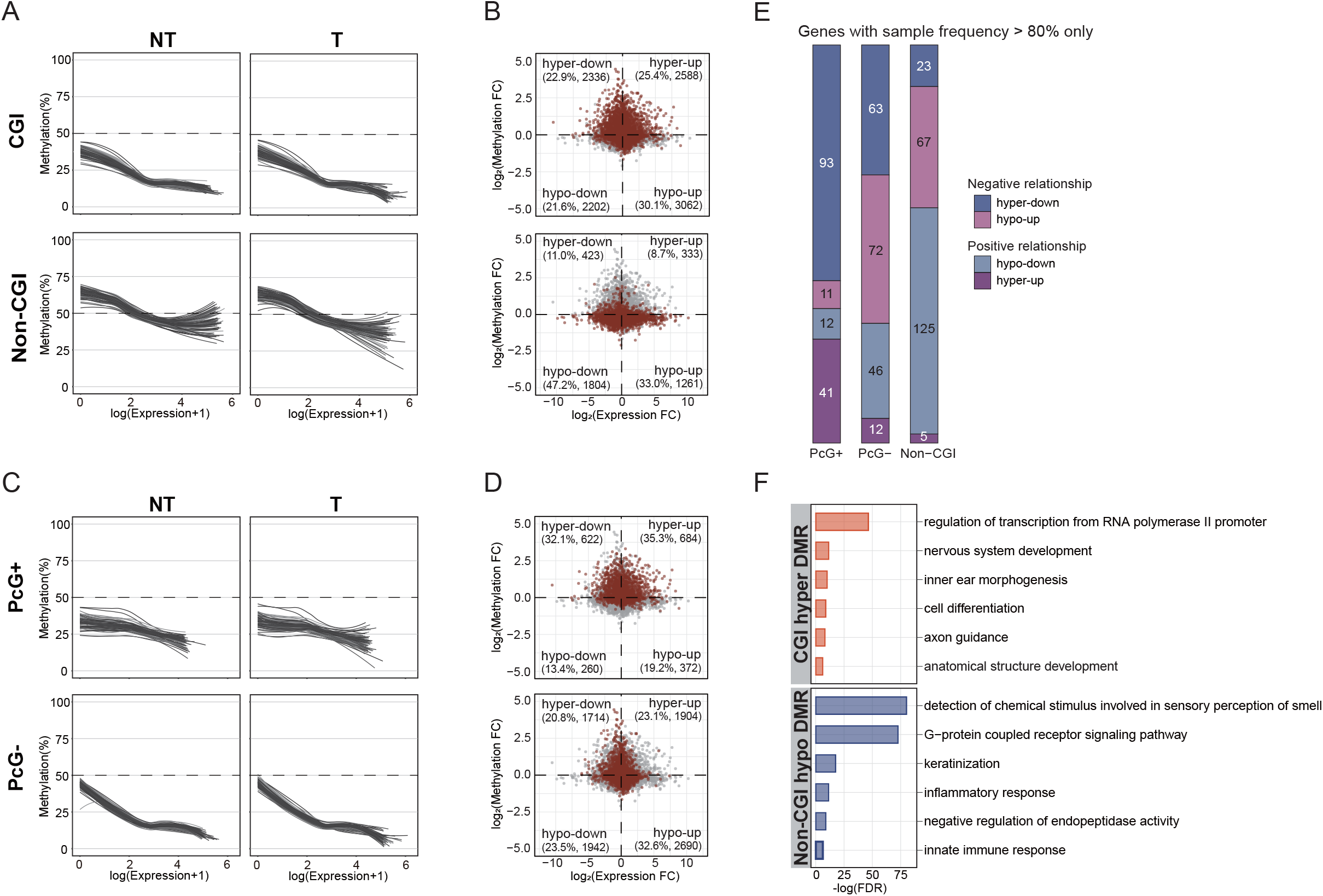
Linking methylation changes to gene expression. **A**. Curves illustrating the relationship between promoter methylation (y-axis) and gene expression (x-axis) for CGI (top) and non-CGI (bottom) promoters in NT and T tissues. Each line represents an individual patient. **B**. Scatter plots illustrating the log_2_FC of average gene expression and DNA methylation across 79 samples. Each quadrant corresponds to one of the four categories: hyper-down (hypermethylation-downregulation), hyper-up (hypermethylation-upregulation), hypo-down (hypomethylation-downregulation), and hypo-up (hypomethylation-upregulation). Gray dots represent all genes, and colored dots highlight genes classified into each group. The top panel corresponds to CGI genes, while the bottom panel represents non-CGI genes. **C**. Curves analogous to A, generated in the PcG context. PcG^+^ on the top, PcG^−^ on the bottom. **D**. The same analysis as in B, PcG^+^ genes on top and PcG^−^ genes below. **E**. Bar plots showing, for PcG^+^, PcG^−^ and non-CGI genes, the proportion of genes that were consistently assigned to the same group (one of hyper-down, hyper-up, hypo-down, or hypo-up) in over 80% of samples (at least 63 out of 79). **F**. Bar plots of the top GO terms for genes with DMRs. Functional terms were assigned by DAVID (https://david.ncifcrf.gov/). The −log (FDR) values are shown in orange for CGI hypermethylated DMRs (top) and in blue for non-CGI hypomethylated DMRs (bottom).

To evaluate whether the presence of PcG marks is associated with the methylation-expression relationship, we performed the same analysis for PcG^+^ and PcG^−^ gene groups, respectively. As shown in Figure 3C, both groups retained the overall inverse methylation–expression trend. However, PcG^+^ promoters displayed weaker and more variable correlations, suggesting that PcG occupancy may buffer or uncouple methylation effects on transcription. Consistent with this, Figure 3D shows that genes with paradoxical hyper-up relationship were more enriched in the PcG^+^ group than in PcG^−^, reinforcing the idea that hypermethylation at PcG^+^ promoters may not always result in transcriptional repression.

These patterns remained consistent when restricting our analysis to genes exhibiting a consistent methylation change (canonical or paradoxical) in > 80 % of patients (Figure 3E). Among PcG^+^ promoters, approximately 85% exhibited hypermethylation, and notably, a substantial fraction (∼70%, 93 hyper-ups/134 hypers) of these showed paradoxical increases in gene expression. In contrast, PcG^−^ promoters showed a more balanced distribution between hypo-up and hyper-dn categories, with paradoxical hyper-up events being rare. For non-CGI promoters, the most frequent consistent pattern was hypomethylation (pink and light-blue), dominated by hypo-dn events, while hypo-up cases were relatively infrequent.

All these findings were further validated in genes with statistically significant changes in both methylation and expression, i.e., DMR-DEG pairs—genes harboring differentially methylated regions (DMRs) and differentially expressed genes (DEGs) (Supple Fwig 1A-F) (Materials and Methods; Supplementary Tables 2-3).

Moreover, gene ontology (GO) analysis of genes with DMR revealed that CGI-hypermethylated genes were strongly enriched for the term “regulation of transcription from RNA polymerase II promoter”, underscoring a predominance of transcription factors (TFs) (Figure 3F). In contrast, non-CGI hypomethylated genes were associated with neuronal signaling and inflammatory response pathways (Figure 3F). Notably, hypermethylated genes in both PcG^+^ and PcG^−^ groups were enriched for TFs, suggesting that promoter hypermethylation preferentially targets TF genes regardless of PcG occupancy (Supple Fwig 1A-F).

### Effect of methylation on the expression of TF genes is varied with PcG^+^- and PcG^−^promoters in tumors

Based on the observation that genes with CGI hypermethylation were predominantly associated with TFs in Figure 3F, we further investigated methylation alterations in tumors, analyzing TFs and non-TFs separately.

Our analysis revealed that a higher proportion of TFs (∼85%) have CGI promoters compared to non-TFs (∼72%) (Figure 4A). By mapping DMRs to gene levels, we examined significant methylation alterations in these two groups during tumorigenesis. Figure 4B shows that while both TFs and non-TFs with CGI promoters are predominantly hypermethylated in tumors, TF hypermethylation is significantly more prevalent than in non-TFs. Specifically, in the CGI category, compared with 68.7% of genes hypermethylated overall, 87% of TFs were hypermethylated, versus 62.9% of non-TFs. Even in the non-CGI category, where hypomethylation predominates (93.4%), TFs still exhibited a significantly higher proportion of hypermethylation than non-TFs; compared with 6.6% of total genes hypermethylated, 26.3% of TFs vs. only 5.8% of non-TFs.

**Figure 4.**
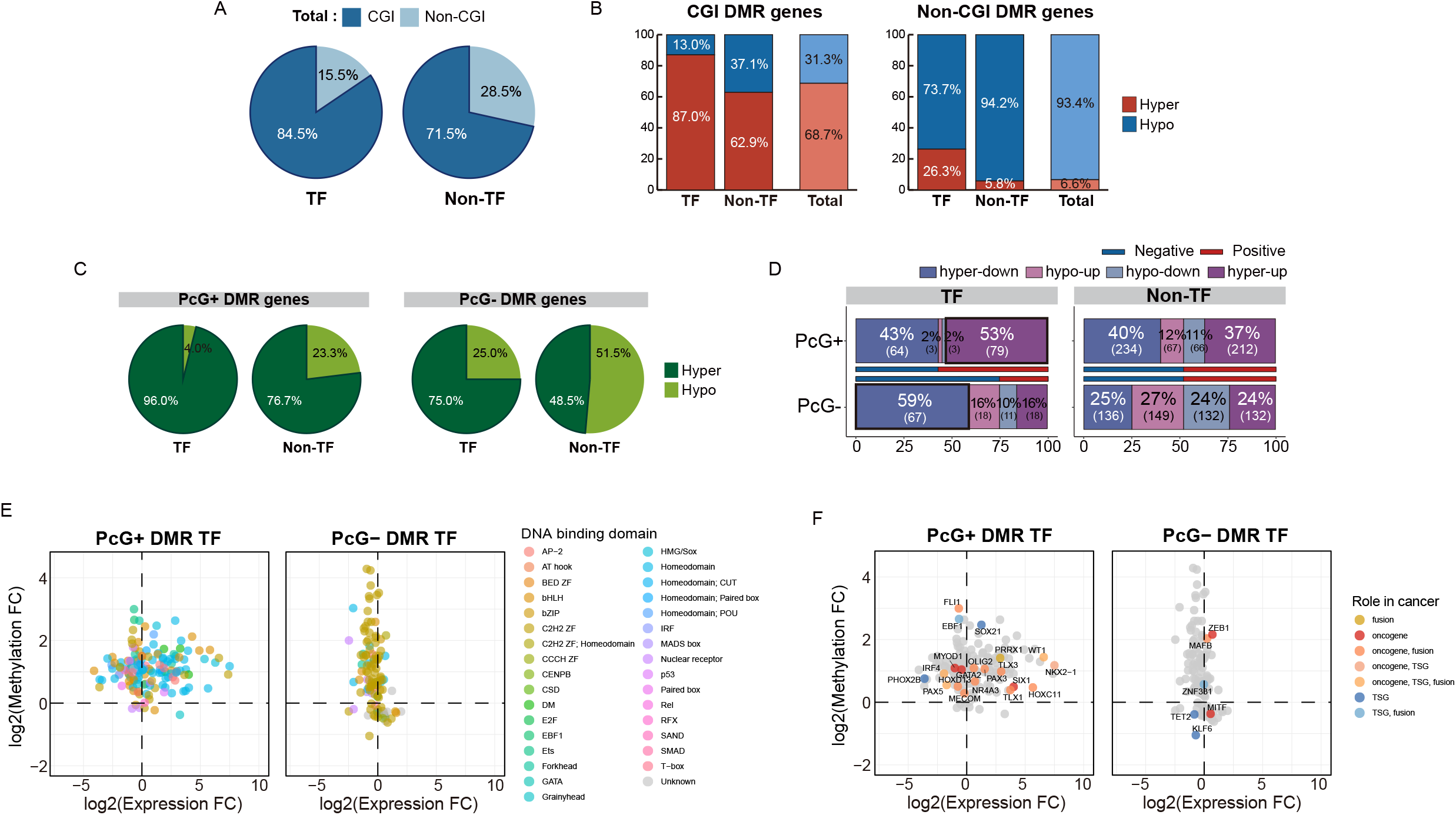
Methylation and gene expression relationships in TFs distinct from those in non-TFs, detected in T compared to NT conditions. **A**. Pie plots showing the ratio of CGI and non-CGI categories in TF genes, and non-TF genes. **B**. Bar graphs comparing the distribution of hypermethylated (red) and hypomethylated (blue) genes in CGI and non-CGI DMR genes. **C**. (Left) Pie charts showing the proportions of hypermethylation in DMRs between PcG^+^ TFs and PcG^+^ non-TFs. (Right) Pie charts displaying the proportions of hypermethylation in DMRs between PcG^−^ TFs and PcG^−^ non-TFs. **D**. Proportions of the four types of methylation and gene expression relationships for TFs and non-TFs in PcG context. **E**. Scatter plots showing the log_2_FC of average gene expression (x-axis) and DNA methylation (y-axis) for CGI DMR TFs. Each point represents a TF, colored according to its DNA-binding domain. DNA-binding domain information was obtained from Lambert et al. (2018) [47] (https://humantfs.ccbr.utoronto.ca/). **F**. Both PcG^+^ and PcG^−^ TFs were matched to genes in the COSMIC database, where classifications as oncogenes or tumor suppressor genes (TSGs) were available.

In the PcG^+^ category, both TFs and non-TFs showed substantial hypermethylation in tumors; however, hypermethylation was notably more prominent in TFs compared to non-TFs (96.0% vs. 76.7%). Similarly, in the PcG^−^ category, TFs still exhibited a clear tendency towards hypermethylation (75.0% in TFs vs. 48.5% in non-TFs), while non-TFs did not show a significant bias in methylation direction (hyper-versus hypomethylation) (Figure 4C).

Next, we examined the relationship between methylation and gene expression for TFs and non-TFs separately. It is important to clarify that when we refer to the relationship between methylation and gene expression for TFs, we specifically mean how methylation in the promoters of TFs affects their own expression, rather than how these TFs regulate the expression of their target genes.

Notably, approximately 43% of DMRs from TFs in the PcG^+^ category were associated with hypermethylation-driven downregulation (hyper-dn, dark-blue), whereas about 53% were linked to the paradoxical hypermethylation-driven upregulation (hyper-up, purple). In contrast, 59% of DMRs from TFs in the PcG^−^ category were associated with hypermethylation-driven downregulation (hyper-dn, dark-blue), while only 16% were classified as hyper-up (purple). These findings suggest that, for TFs, hypermethylation in PcG^+^ TFs is paradoxically more likely associated with upregulation, whereas hypermethylation in PcG^−^ TFs tends to be canonically linked to suppressed expression. On the other hand, hypermethylation in PcG^+^ non-TF genes appeared to have a slightly greater canonical negative impact compared to a paradoxical positive impact (40% hyper-dn, dark blue vs. 37% hyper-up, purple). Meanwhile, PcG^−^ non-TFs were more evenly distributed across the four categories of methylation–gene expression relationships, showing a balanced pattern of association (Figure 4D).

Individual gene analysis of CGI DMR TFs in Figure 4E further confirmed the frequent paradoxical upregulation of hypermethylated PcG^+^ TFs, consistent with the trends observed in Figure 4D. Interestingly, domain analysis revealed that PcG^+^ TFs were enriched for homeodomains, whereas PcG^−^ TFs predominantly contained zinc finger domains (Figure 4E). Moreover, PcG^+^ TFs showed greater overlap with oncogenes listed in the COSMIC database (Figure 4F), suggesting that upregulation of homeodomain-containing TFs with PcG^+^ promoters may play a key role in driving tumorigenesis.

## Discussion

Understanding the interplay between DNA methylation and gene expression is fundamental in cancer epigenetics. In this study, we leveraged paired non-tumor–tumor samples from the same individuals to explore how methylation dynamics correlate with gene expression at an individual level. This patient-matched approach enabled us to uncover significant heterogeneity that average-based comparisons would otherwise conceal, providing key insights into TF regulation in tumors—especially in Polycomb-marked (PcG^+^) genes.

One of our central findings is that methylation changes do not predict gene expression changes uniformly across individuals. While we confirmed the classical trends of CGI hypermethylation and non-CGI hypomethylation in tumors—consistent with earlier work [29-32]—our paired analyses revealed substantial inter-individual variability in how these methylation changes affect gene expression. For example, contrary to the conventional expectation that hypomethylation leads to gene activation, we found that non-CGI hypomethylation can be linked to decreased gene expression (Figure 3E). This suggests that baseline methylation levels of non-CGI promoters in non-tumor tissues may establish a particular chromatin state that persists despite hypomethylation. These promoters may require additional regulatory inputs—such as TF binding or enhancer interactions—to fully activate transcription.

When stratifying CGI genes into PcG^+^ vs. PcG^−^ categories, we observed that PcG^−^ genes tend to be more strongly repressed upon hypermethylation (Figure 3C), consistent with conventional models. By contrast, PcG^+^ genes showed relative resistance to methylation-induced silencing, suggesting that PcG occupancy confers epigenetic plasticity in tumors. This observation aligns with previous studies showing that PcG^+^ promoters can remain poised for activation, especially in response to specific developmental or environmental signals [21, 33, 34]. One of the most intriguing outcomes of our work is the paradoxical hypermethylation coupled with transcriptional upregulation of TF genes that carry PcG marks in their promoters. This challenges the established view that promoter methylation always enforces silencing, indicating that TF expression in tumors is governed by additional regulatory mechanisms beyond DNA methylation alone.

Among PcG^+^ promoters, TF genes were substantially more likely than non-TF genes to maintain transcriptional activity despite hypermethylation (Figure 4D). Given that TFs orchestrate gene regulatory networks, their expression is likely controlled by complex epigenetic crosstalk, including histone modifications and higher-order chromatin organization [35, 36]. This finding is particularly relevant in cancer biology, suggesting that tumors preserve or enhance the expression of certain TFs—despite widespread epigenetic reprogramming.

One plausible explanation for hypermethylation plus upregulation in PcG^+^ TF promoters is that distal enhancer activity counteracts the silencing effect of promoter hypermethylation. Indeed, in CRC, Zheng et al. (2021) showed that many PcG^+^ CGI promoters become hypermethylated yet remain transcriptionally active, driven by active distal enhancers that recruit TFs and coincide with local loss of H3K27me3 marks [22].

In our study, we extend these observations specifically to TFs themselves. We find that PcG^+^ TFs often exhibit promoter hypermethylation in tumors yet remain transcriptionally upregulated, whereas TFs that lack PcG marks in their promoters tend not to be upregulated under hypermethylated conditions. This suggests that PcG occupancy confers a unique epigenetic plasticity, allowing TF promoters to stay active—even when hypermethylated—in the context of tumor-specific enhancer signals. Consequently, these upregulated TFs may drive the expression of downstream cancer-associated genes, further supporting oncogenic programs.

Meanwhile, single-cell epigenomic approaches increasingly reveal heterogeneous methylation landscapes in tumors, indicating that not all tumor cells adhere to uniform epigenetic patterns [37, 38]. However, many such studies lack paired methylation and transcription data at the single-patient level. Our work bridges this gap by providing direct evidence, across multiple individuals, that PcG^+^ TF promoters can evade hypermethylation-mediated repression. This underscores unrecognized complexity in tumor gene regulation.

Still, key questions remain: (1) What specific mechanisms enable PcG^+^ TF promoters to bypass methylation repression, (2) Are these findings applicable across diverse cancer types, and (3) How might these observations inform cancer therapy? Future studies should integrate chromatin accessibility (ATAC-seq) and histone modification data to delineate the interplay between DNA methylation, enhancer activity, and transcriptional output. Although our study focused on CRC, performing similar analyses in other malignancies will reveal whether PcG^+^ TF hypermethylation plus upregulation is a widespread tumor adaptation. If PcG^+^ TFs selectively escape methylation repression, they may be promising therapeutic targets or biomarkers in tumors that leverage epigenetic plasticity to sustain oncogenic programs.

## Conclusion

In sum, our study challenges the simplistic view of DNA methylation as a straightforward on– off switch for gene expression. By specifically analyzing paired non-tumor–tumor samples, we reveal notable inter-individual variability in the relationship between promoter methylation and transcription, highlighting a striking epigenetic paradox in PcG^+^ TF regulation. These insights expand our understanding of tumor epigenetics, emphasizing the importance of considering chromatin context—particularly Polycomb-associated states—in cancer. Further investigations into PcG^+^ TF regulation could offer new avenues for epigenetic-based therapies, targeting the mechanisms that allow tumors to circumvent methylation-mediated repression yet sustain high-level TF activity.

## Materials and Methods

### Obtaining TBS and RNA-seq data

We obtained TBS (K-BDS, KAP220182) and RNA-seq data (EGA, EGAD00001006985), which deposited data produced from a total of 80 Korean CRC patient samples excluding TNM stage 4 samples. Refer to our previous paper [28] for the information on samples.

### Analysis of methylome data

Raw reads generated by TBS were processed using ‘FastQC’ (v0.11.9) [39] for quality checks and ‘Trim Galore’ (v0.6.7) (https://www.bioinformatics.babraham.ac.uk/projects/trim_galore/) [40] for adaptor trimming. Subsequently, after indexing of reference genome (GRCh38.p12) using ‘Bismark’(v0.22.1) [41], the cleaned reads were mapped to it. Additionally, cytosine report files containing CpG methylation context were obtained using the ‘Bismark methylation extractor’. The beta values of each region were calculated using ‘percMethylation’ function of the ‘MethylKit’ (v1.17.5) of R package. DMRs were identified using the ‘methylKit’ (v1.17.5) [42] of R package with the ‘region resolution’ option. Promoter regions were defined as TSS 1500∼2000 using the GTF file of GENCODE GRCh38.p12, along with the ‘Detailed Annotation’ output of annotatePeaks.pl from ‘Homer’ (v4.11.1) [43]. After normalization, DMRs were identified by two thresholds, |methdiff| ≥ 10, Q-value < 0.01. Genes that exhibited both hypermethylation and hypomethylation simultaneously were excluded.

### Analysis of RNAs-seq data

After quality checks using ‘FastQC’(v0.11.9) [39], the reference genome (GRCh38.p12) was indexed with ‘STAR’ (v020201) [44], to which raw read were mapped. Subsequently, the raw read count table was generated using ‘htseq-count’ from ‘HTSeq’ (v0.11.2) [45]. Gene annotation was performed with the GTF file of GENCODE GRCh38.p12. Normalization of raw read counts and identification of DEGs were carried out using ‘DESeq2’(v1.32.0) [46], with thresholds of |log_2_FC|≥1, Q-value < 0.01. DMR-DEG pairs were defined by matching genes with DMRs to those with DEGs, and further filtered to retain only pairs showing a significant cross-sample correlation (|Pearson correlation coefficient| > 0.4, P < 0.01).

### Filtering criteria for 14,042 protein-coding genes

We obtained expression values for a total of 58,721 genes via RNA-seq and % methylation values for a total of 46,390 genes via TBS. Among them, we selected genes with both expression and methylation data in the promoter region, and then selected only protein-coding genes to obtain a total of 14,042 genes.

### Defining gene groups: CGI and non-CGI genes, and PcG^+^ and PcG^−^-CGI genes

To classify genes into CGI and non-CGI groups, we used the ‘readFeatureFlank’ function from the ‘MethylKit’ package, utilizing CGI information for gene promoters from the GRCh38.cpgisland.bed file, obtained from the UCSC genome browser (https://genome.ucsc.edu/cgi-bin/hgTables). For the classification of PcG^+^ and PcG^−^ groups, we used the ChromHMM annotation file (ENCSR807KHQ), which contains information on 18 chromatin states derived from colonic mucosa, obtained from the ENCODE project (https://www.encodeproject.org/). Based on the ChromHMM 18-state model, we then overlapped the CGI promoter regions defined earlier with PcG-related regions (‘ReprPC’ and ‘ReprPCWk’). Genes overlapping these regions were classified as PcG^+^-CGI genes, while genes without overlap were classified as PcG^−^-CGI genes.

### Obtaining TF gene list

We downloaded the official list of human TFs from the website (https://humantfs.ccbr.utoronto.ca/) from Lambert et al. (2018) [47].

### Statistical analysis and visualization

All statistical analyses were performed with R (v4.1.0) (https://www.R-project.org/), using ggplot2(v3.3.4) [48] for visualization. GO analysis was conducted with DAVID (ver. 6.8) [49, 50] (https://david.ncifcrf.gov/), where terms in the biological process were selected based on FDR. A correlation test between methylation levels and gene expression levels of DMR-DEG pairs was carried out with the ‘cor’ function in R.

## Ethics approval and consent to participate

Not applicable

## Abbreviations

CGI: CpG island
CRC: Colorectal cancer
DEG: Differentially expressed gene
DMR: Differentially methylated region
FC: Fold change
Non-CGI: Non CpG island
NT: Non-tumor
PcG: Polycomb group
T: Tumor
TBS: Targeted bisulfite sequencing
TF: Transcription Factor

## Data availability

Raw sequencing reads of RNA-seq data can be downloaded from the European Genome-phenome Archive (EGA) with the data accession number EGAD00001006985 (under study number EGAS00001005068) and TBS data can be downloaded from Korea BioData Station (K-BDS) with the BioProject ID KAP220182.

## Author contributions

SSC & SH supervised the research and wrote the manuscript. MKK, GP conducted the analysis and also contributed to writing the manuscript. DP & SH participated in discussion related to study design and assisted in the writing process.

## Additional information

**Correspondence** should be addressed to SSC.

## References

1. Moore, L.D., T. Le, and G.P. Fan, DNA Methylation and Its Basic Function. Neuropsychopharmacology, 2013. 38(1): p. 23–38.

2. Smith, Z.D. and A. Meissner, DNA methylation: roles in mammalian development. Nature Reviews Genetics, 2013. 14(3): p. 204–220.

3. Deaton, A.M. and A. Bird, CpG islands and the regulation of transcription. Genes Dev, 2011. 25(10): p. 1010–22.

4. Héberlé, É. and A.F. Bardet, Sensitivity of transcription factors to DNA methylation. DNA Methylation, 2019. 63(6): p. 727–741.

5. Jones, P.L., et al., Methylated DNA and MeCP2 recruit histone deacetylase to repress transcription. Nat Genet, 1998. 19(2): p. 187–91.

6. Fuks, F., DNA methylation and histone modifications: teaming up to silence genes. Curr Opin Genet Dev, 2005. 15(5): p. 490–5.

7. Wade, P.A., et al., Mi-2 complex couples DNA methylation to chromatin remodelling and histone deacetylation. Nat Genet, 1999. 23(1): p. 62–6.

8. Robertson, K.D., DNA methylation and human disease. Nat Rev Genet, 2005. 6(8): p. 597–610.

9. Fleisher, A.S., et al., Hypermethylation of the hMLH1 gene promoter is associated with microsatellite instability in early human gastric neoplasia. Oncogene, 2001. 20(3): p. 329–35.

10. Catteau, A. and J.R. Morris, BRCA1 methylation: a significant role in tumour development? Semin Cancer Biol, 2002. 12(5): p. 359–371.

11. Kondo, Y., L. Shen, and J.P. Issa, Critical role of histone methylation in tumor suppressor gene silencing in colorectal cancer. Mol Cell Biol, 2003. 23(1): p. 206–15.

12. Yoshiura, K., et al., Silencing of the E-cadherin invasion-suppressor gene by CpG methylation in human carcinomas. Proc Natl Acad Sci U S A, 1995. 92(16): p. 7416–9.

13. Kim, J.S., et al., Aberrant methylation of H-cadherin (CDH13) promoter is associated with tumor progression in primary nonsmall cell lung carcinoma. Cancer, 2005. 104(9): p. 1825–33.

14. Sheaffer, K.L., E.N. Elliott, and K.H. Kaestner, DNA Hypomethylation Contributes to Genomic Instability and Intestinal Cancer Initiation. Cancer Prev Res (Phila), 2016. 9(7): p. 534–46.

15. Ehrlich, M., DNA hypomethylation in cancer cells. Epigenomics, 2009. 1(2): p. 239–59.

16. Lakshminarasimhan, R. and G.N. Liang, The Role of DNA Methylation in Cancer. DNA Methyltransferases - Role and Function, 2016. 945: p. 151–172.

17. de Mendoza, A., et al., Large-scale manipulation of promoter DNA methylation reveals context-specific transcriptional responses and stability. Genome Biology, 2022. 23(1).

18. Rauluseviciute, I., F. Drablos, and M.B. Rye, DNA hypermethylation associated with upregulated gene expression in prostate cancer demonstrates the diversity of epigenetic regulation. Bmc Medical Genomics, 2020. 13(1).

19. Parreno, V., A.M. Martinez, and G. Cavalli, Mechanisms of Polycomb group protein function in cancer. Cell Research, 2022. 32(3): p. 231–253.

20. Schlesinger, Y., et al., Polycomb-mediated methylation on Lys27 of histone H3 pre-marks genes for de novo methylation in cancer. Nat Genet, 2007. 39(2): p. 232–6.

21. Hahn, M.A., et al., Loss of the polycomb mark from bivalent promoters leads to activation of cancer-promoting genes in colorectal tumors. Cancer Res, 2014. 74(13): p. 3617–3629.

22. Zheng, Y., et al., A pan-cancer analysis of CpG Island gene regulation reveals extensive plasticity within Polycomb target genes. Nat Commun, 2021. 12(1): p. 2485.

23. Pherson, M., et al., Polycomb repressive complex 1 modifies transcription of active genes. Sci Adv, 2017. 3(8): p. e1700944.

24. Parreno, V., et al., Transient loss of Polycomb components induces an epigenetic cancer fate. Nature, 2024. 629(8012): p. 688–696.

25. Yin, Y.M., et al., Impact of cytosine methylation on DNA binding specificities of human transcription factors. Science, 2017. 356(6337).

26. Hoadley, K.A., et al., Cell-of-Origin Patterns Dominate the Molecular Classification of 10,000 Tumors from 33 Types of Cancer. Cell, 2018. 173(2): p. 291–304 e6.

27. Toyota, M., et al., CpG island methylator phenotype in colorectal cancer. Proc Natl Acad Sci U S A, 1999. 96(15): p. 8681–6.

28. Kim, J., et al., Transcriptomes of the tumor-adjacent normal tissues are more informative than tumors in predicting recurrence in colorectal cancer patients. J Transl Med, 2023. 21(1): p. 209.

29. Figueiredo, J.C., et al., Global DNA hypomethylation (LINE-1) in the normal colon and lifestyle characteristics and dietary and genetic factors. Cancer Epidemiol Biomarkers Prev, 2009. 18(4): p. 1041–9.

30. Zelic, R., et al., Global Hypomethylation (LINE-1) and Gene-Specific Hypermethylation (GSTP1) on Initial Negative Prostate Biopsy as Markers of Prostate Cancer on a Rebiopsy. Clin Cancer Res, 2016. 22(4): p. 984–92.

31. Sproul, D. and R.R. Meehan, Genomic insights into cancer-associated aberrant CpG island hypermethylation. Brief Funct Genomics, 2013. 12(3): p. 174–90.

32. Moarii, M., et al., Changes in correlation between promoter methylation and gene expression in cancer. BMC Genomics, 2015. 16: p. 873.

33. Ferrari, K.J., et al., Polycomb-dependent H3K27me1 and H3K27me2 regulate active transcription and enhancer fidelity. Mol Cell, 2014. 53(1): p. 49–62.

34. Cruz-Molina, S., et al., PRC2 Facilitates the Regulatory Topology Required for Poised Enhancer Function during Pluripotent Stem Cell Differentiation. Cell Stem Cell, 2017. 20(5): p. 689–705 e9.

35. Lee, T.I., et al., Control of developmental regulators by Polycomb in human embryonic stem cells. Cell, 2006. 125(2): p. 301–13.

36. Xie, W., et al., Epigenomic analysis of multilineage differentiation of human embryonic stem cells. Cell, 2013. 153(5): p. 1134–48.

37. Li, S., et al., Chromatin accessibility dynamics in colorectal cancer liver metastasis: Uncovering the liver tropism at single cell resolution. Pharmacol Res, 2023. 195: p. 106896.

38. Terekhanova, N.V., et al., Epigenetic regulation during cancer transitions across 11 tumour types. Nature, 2023. 623(7986): p. 432–441.

39. FastQC. 2015.

40. Martin, M., Cutadapt removes adapter sequences from high-throughput sequencing reads. 2011, 2011. 17(1): p. 3.

41. Krueger, F. and S.R. Andrews, Bismark: a flexible aligner and methylation caller for Bisulfite-Seq applications. Bioinformatics, 2011. 27(11): p. 1571–2.

42. Akalin, A., et al., methylKit: a comprehensive R package for the analysis of genome-wide DNA methylation profiles. Genome Biol, 2012. 13(10): p. R87.

43. Heinz, S., et al., Simple combinations of lineage-determining transcription factors prime cis-regulatory elements required for macrophage and B cell identities. Mol Cell, 2010. 38(4): p. 576–89.

44. Dobin, A., et al., STAR: ultrafast universal RNA-seq aligner. Bioinformatics, 2013. 29(1): p. 15–21.

45. Anders, S., P.T. Pyl, and W. Huber, HTSeq--a Python framework to work with high-throughput sequencing data. Bioinformatics, 2015. 31(2): p. 166–9.

46. Love, M.I., W. Huber, and S. Anders, Moderated estimation of fold change and dispersion for RNA-seq data with DESeq2. Genome Biol, 2014. 15(12): p. 550.

47. Lambert, S.A., et al., The Human Transcription Factors. Cell, 2018. 172(4): p. 650–665.

48. Wickham, H., ggplot2: Elegant Graphics for Data Analysis. 2016: Springer-Verlag New York.

49. Sherman, B.T., et al., DAVID: a web server for functional enrichment analysis and functional annotation of gene lists (2021 update). Nucleic Acids Res, 2022. 50(W1): p. W216–W221.

50. Huang da, W., B.T. Sherman, and R.A. Lempicki, Systematic and integrative analysis of large gene lists using DAVID bioinformatics resources. Nat Protoc, 2009. 4(1): p. 44–57.

51. Liska, O., et al., TFLink: an integrated gateway to access transcription factor-target gene interactions for multiple species. Database-the Journal of Biological Databases and Curation, 2022. 2022.

