## supplementary figure for "Transcription factors overcome the repressive impact of Polycomb-associated methylation in tumors": 20250501_supple_figure.docx

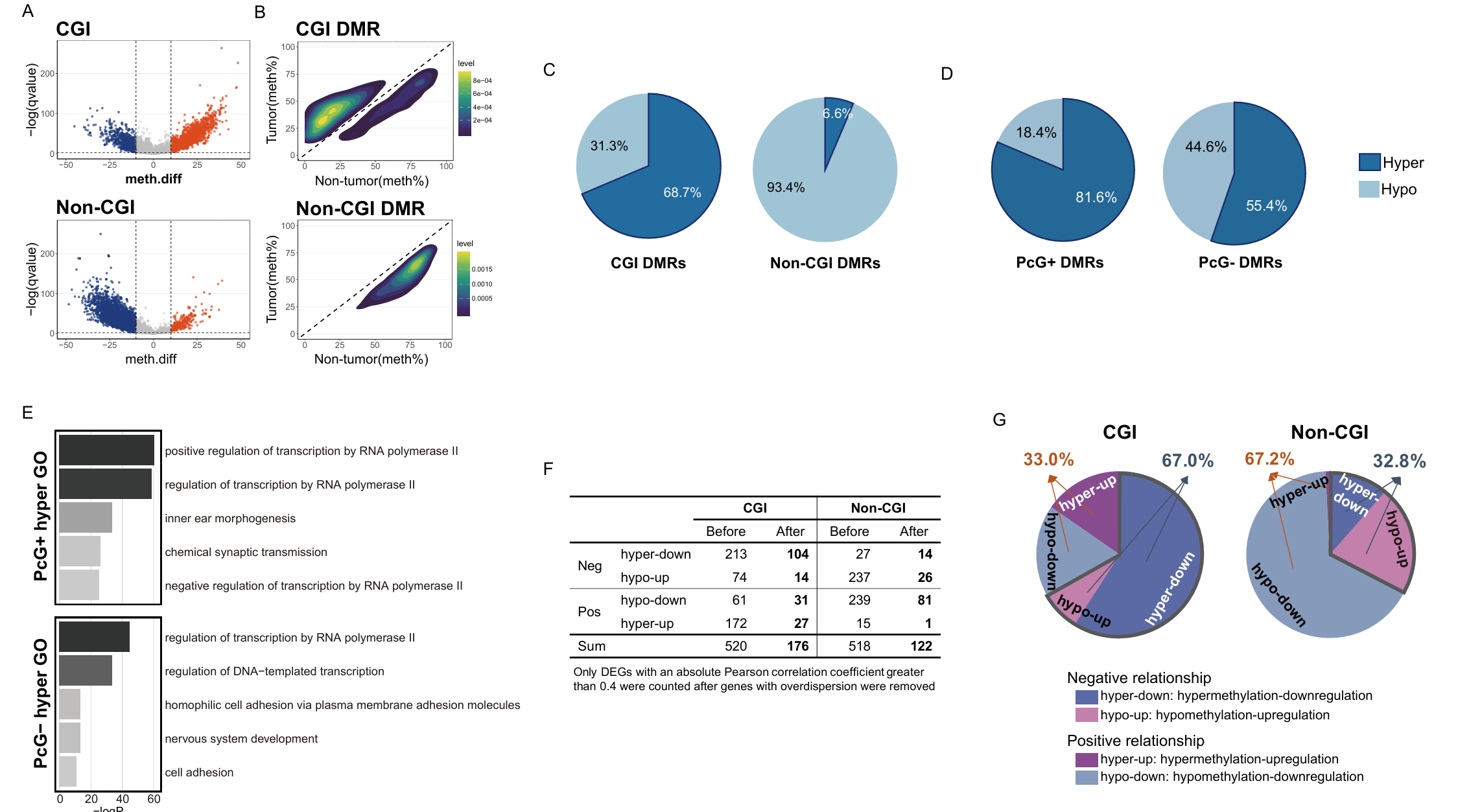

**Supplementary Figure 1. A.** Volcano plots of % methylation for genes with CGI and non-CGI promoters. DMRs were identified with Q-value < 0.01 and |meth.diff| ≥ 10, indicated as hypermethylated (orange) and hypomethylated (blue) in the plot. Non-significant DMRs are shown in gray. **B.** Density plots of DMRs identified from genes with CGI and non-CGI promoters, showing methylation percentages in tumor and non-tumor samples. The density increases as colors transition from blue to yellow. The dotted diagonal line represents equal methylation in tumor and non-tumor samples, with points above the line being hypermethylated and below the line being hypomethylated in tumor samples compared to non-tumor samples. **C.** Pie charts showing the proportions of hypermethylation (dark blue) and hypomethylation (light-blue) in the DMRs from CGI promoters (CGI DMRs) and non-CGI promoters (non-CGI DMRs). **D.** Pie charts similar to C, shown separately for PcG⁺ (left) and PcG⁻ (right) groups. **E.** Bar plots of top GO terms for genes associated with hypermethylated DMRs in PcG^+^ and PcG^–^ categories. **F.** Table showing the numbers of two types of DMR-DEG pairs before and after applying a filter for significant correlation (|Pearson correlation coefficient| > 0.4, P < 0.01). **G.** Pie charts showing the proportions of each subgroup obtained from the DMR-DEG pair relationships. Two types of relationships in DMR-DEG pairs are indicated: a negative relationship (hyper-down and hypo-up) with a bold border, and a positive relationship (hyper-up and hypo-dn) with no border line.

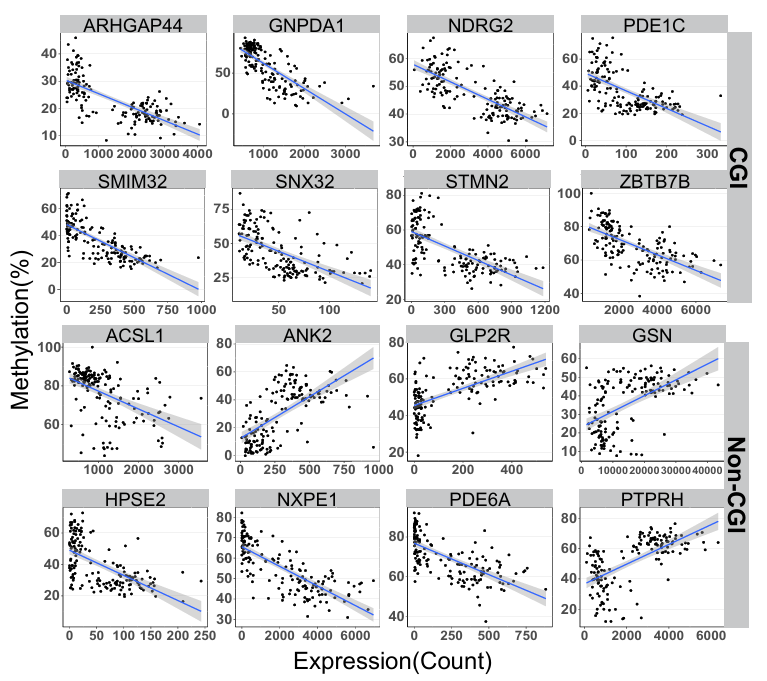

**Supplementary Figure 2.** Correlation plots showing DMR-DEG pairs with a significant correlation between methylation and gene expression. DMR-DEG pairs was selected with |Pearson correlation coefficient| > 0.4 and P-value < 0.01.
